# Photodynamic Inactivation reduces the diversity and changes the composition of bacterial and fungal communities associated with leaf surfaces

**DOI:** 10.1101/2021.04.12.439450

**Authors:** Robert R. Junker, Lisa-Maria Ohler, Christoph Hamminger, Kristjan Plaetzer

## Abstract

Plant surfaces are colonized by a myriad of microorganisms including mutualistic strains and pathogens. Particularly in agricultural systems applications are required that protect the plants against pathogens without negative side effects on the environment and humans. Photodynamic Inactivation (PDI) has been demonstrated to be a promising approach to efficiently fight plant pathogens. Based on its mechanism of action, the light-induced and photosensitizer-mediated overproduction of reactive oxygen species in target cells, PDI is likely to generally inactivates microorganisms on plants irrespective of their pathogenicity. In order to prove this hypothesis we used next-generation 16S rRNA gene amplicon sequencing to characterize the bacterial and fungal communities associated with leaf surfaces of *Arabidopsis thaliana* before and after the photodynamic treatment using the chlorine e6 derivative B17-0024 as photoactive compound and showed that this treatment reduced the microbial richness and altered the microbial community composition. These findings may help to develop effective pathogen-control strategies and may also stimulate research on plant-microbe interactions exploiting the potential of PDI.

## Introduction

Leaf surfaces are one of the largest habitats for bacteria and fungi that together with further microorganisms comprise the plant microbiome [1]. Many of these microorganisms are plant pathogens that cause dramatic diseases in both natural and agricultural systems. The global crop losses caused by bacterial and fungal plant pathogens are estimated to account for 16% of the overall agricultural yield [2]. Additional to pathogens that harm the plants they are infecting, many epiphytic microorganisms help the plants to resist abiotic stresses, and protect the plants against diseases and herbivores [3]. These beneficial effects result from the presence of individual strains that, for example, can suppress disease symptoms and reduce the growth of pathogens [4]. Alternatively, a diverse microbiome, i.e. an assemblage of microorganisms that individually may not have a direct effect on plants, is strongly contributing to plant healthy and growth. For instance, the diversity of epiphytic bacteria was negatively correlated to the severity of the southern leaf blight disease in maize [5] indicating that pathogens are less likely to establish on plant surfaces that are associated with species-rich microbial communities [6]. Accordingly, it is expected that the beneficial functions of the plant microbiome will significantly contribute to eco-friendly agriculture and thus economic growth and are therefore increasingly studied in academia and industry [7].

Plant pathogens have the potential to strongly reduce crop yield and represent a major threat to food security. The use of pesticides in industrialized monocultural farming is the standard treatment for farmers to combat phytopathogens, but those chemicals have the potential to leach into soil and water and cause pollution. Particularly the application of antibiotics has been criticized for negative side effects on the environment and also the consumers and for being not efficient for a long time due to the development of antibiotic resistance [8]. Recently, Photodynamic Inactivation (PDI) has been proposed as an environmentally-friendly alternative for conventional antibiotics [9]. The mode of action of PDI is based on the combination of two *per se* harmless components. As the first step of the treatment protocol, a photoactive compound, the photosensitizer (PS) is applied for example as aqueous spray and accumulates by various mechanisms in or at the microorganism’s cell wall. Illumination with visible light of an appropriate wavelength (e.g. sunlight) represents the second step and leads to photocatalytic overproduction of reactive oxygen species in microorganisms, which oxidize cellular targets and induce cell death [10]. Another promising approach to reduce the impact of pathogens on crop yield are biocontrol strategies that aim at modulating plant defense mechanisms by the application of individual strains or by engineering beneficial microbiomes [11]. Thus, efficient plant protection measures need to compromise between an effective elimination of the pathogen and the maintenance of a healthy and diverse microbiome. Therefore, particularly for non-targeted approaches that combat non-specifically bacteria and fungi, such as PDI, it is mandatory to evaluate the effects of these measures on the whole plant microbiome. In order to address this open question, we used next-generation 16S rRNA gene amplicon sequencing to characterize the bacterial and fungal communities associated with leaf surfaces of *Arabidopsis thaliana* before and after the photodymanic treatment using the chlorine e6 derivative B17-0024 as photoactive compounds that has been shown to be efficient in removing pathogens from plant surfaces [9]. We hope that our data will help to develop strategies for the application of PDI that efficiently fights pathogens while maintaining beneficial strains and a healthy microbiome.

## Material and Methods

### Plant Material

*Arabidopsis thaliana* plants (Col-0) were cultivated in pots with non-sterile soil inside the greenhouse in the botanical garden at the University of Salzburg until they reached anthesis. Pots and space between soil and leaves was covered with aluminum foil to prevent reintroduction of microorganisms from the soil to the leaves after treatment.

### Photodynamic Inactivation of plant surfaces

*A. thaliana* plants were randomly assigned to either control (*n* = 4) or treatment group (*n* = 4). The plants assigned for treatment were sprayed using a spray bottle containing a solution of 100 µM of B17-0024 (B17-0024 Ce6 15, 17, DMAE; Suncor AgroScience, Mississauga Ontario, Canada) in autoclaved tap water until run-off. Control plants were sprayed with autoclaved tap water without PS [10]. After spraying, the plants were incubated for five minutes in the dark at 26 °C. Illumination was performed from above (average distance between the light source and the plants was 10 cm) using a LED array [12] containing 432 LEDs with a dominant wavelength of 395 nm (L-7113UVC, Kingbright Electronic Europe GmbH, Issum Germany) using an irradiance of 14.8 mW cm^-2^ for 120 minutes thus resulting in a radiant exposure of 106.6 J cm^-2^. Immediately after the treatment, plants were transferred to sterile polypropylene containers with permeable filter lids for enabling gas exchange (Microbox by Sac O2, Deinze, Belgium) to avoid contamination with microorganisms from the environment. For assessing the number of viable bacterial and fungal cells present on the leaf surface, one fully developed, intact leaf of each plant was removed under sterile conditions and pressed on agar plates with non-selective growth medium for bacteria (lysogeny broth) and fungi (malt extract). This procedure was repeated prior to treatment (pre) immediately after treatment (d0) and three days after treatment (d3).

### Next-generation 16S rRNA gene amplicon sequencing

For microbial DNA extraction, leaves were harvested from *A. thaliana* plants using sterile forceps prior to treatment (pre) immediately after treatment (d0) and three days after treatment (d3). Each biological replicate was collected from a different plant. Leaves were placed in BashingBeads Lysis tubes from the Xpedition™ Fungal/Bacterial DNA Miniprep kit (Zymo Research) containing lysis solution. Lysis tubes were sonicated for 7 min to detach microorganisms from plant surfaces and the plant material was subsequently removed from the solution. Lysis tubes were shaken 5 min at 16 Hz using a ball mill to extract and purify DNA following the manufacturer’s guidelines (Zymo Research). Microbiome profiling of isolated DNA samples was performed by Eurofins Genomics (Ebersberg, Germany). Eurofins Genomics amplified and Illumina MiSeq sequenced the V3-V4 region of the 16S rRNA gene to identify bacterial operational taxonomic units (OTUs) and the ITS2 region for fungal OTUs following the standard procedure “InView - Microbiome Profiling 3.0 with MiSeq”. Sequences were demultiplexed, the primers were clipped, forward and reverse reads were merged and merged reads were quality filtered. Microbiome analysis was performed by Eurofins Genomics using the company’s standard procedure (the following description of analysis is provided by Eurofins Genomics): reads with ambiguous bases (“N”) were removed. Chimeric reads were identified and removed based on the de-novo algorithm of UCHIME [13] as implemented in the VSEARCH package [14]. The remaining set of high-quality reads was processed using minimum entropy decomposition [15,16]. Minimum Entropy Decomposition (MED) provides a computationally efficient means to partition marker gene datasets into OTUs (Operational Taxonomic Units). Each OTU represents a distinct cluster with significant sequence divergence to any other cluster. By employing Shannon entropy, MED uses only the information-rich nucleotide positions across reads and iteratively partitions large datasets while omitting stochastic variation. The MED procedure outperforms classical, identity-based clustering algorithms. Sequences can be partitioned based on relevant single nucleotide differences without being susceptible to random sequencing errors. This allows a decomposition of sequence data sets with a single nucleotide resolution. Furthermore, the MED procedure identifies and filters random “noise” in the dataset, i.e. sequences with a very low abundance (less than 0.02% of the average sample size). To assign taxonomic information to each OTU, DC-MEGABLAST alignments of cluster representative sequences to the sequence database were performed (Reference database: NCBI_nt (Release 2018-07-07)). A most specific taxonomic assignment for each OTU was then transferred from the set of best-matching reference sequences (lowest common taxonomic unit of all best hits). Hereby, a sequence identity of 70% accross at least 80% of the representative sequence was a minimal requirement for considering reference sequences. Further processing of OTUs and taxonomic assignments was performed using the QIIME software package (version 1.9.1, http://qiime.org/) [17]. Abundances of bacterial and fungal taxonomic units were normalized using lineage-specific copy numbers of the relevant marker genes to improve estimates [18].

### Statistical analysis

Prior to the statistical analysis of bacterial and fungal communities associated with leaf surfaces, we performed a cumulative sum scaling (CSS) normalization (R package *metagenomeSeq* v1.28.2) on the count data. Bacterial and fungal richness (i.e. the number of OTUs per sample) was quantified using the R package *vegan* [19]. We performed Kruskal-Wallis rank sum tests to test for significant differences between control and treatment plants. In order to test for significant differences in bacterial and fungal community composition between treatment groups, we performed a Bray-Curtis distance-based redundancy analysis (R package *vegan*) followed by a permutation test. We used Venn diagrams to visualize the number of OTUs unique to or shared by the treatment groups using the R package *VennDiagram*. Additionally, we performed Kruskal-Wallis rank sum tests to test for differences in the abundance of individual strains as a function of treatment groups.

## Results

Both bacterial and fungal communities were strongly affected by PDI treatment. The number of detectable bacterial OTUs dropped from nearly 100 (median value) to 35 directly after the treatment. Three days later, we detected 73 bacterial OTUs on control leaf surfaces, whereas OTU number was still reduced in treated plants (35, Kruskal-Wallis rank sum test: *X*^*2*^ = 11.03, *df* = 3, *p* = 0.012, Fig. 1a). Overall, fungal communities comprised a lower OTU richness with 5 and 3 OTUs (median values) in control plants on day 0 and day 3, respectively. In three out of four samples, we did not detect any fungal OTUs after PDI treatment, one sample featured three OTUs. Three days after the treatment, the number of fungal OTUs was still reduced in two samples, whereas two further samples showed a strong increase in fungal richness with up to 14 OTUs. This clear trend was however not statistically significant (Kruskal-Wallis rank sum test: *X*^*2*^ = 4.75, *df* = 3, *p* = 0.191, Fig. 1b).

**Fig. 1.**
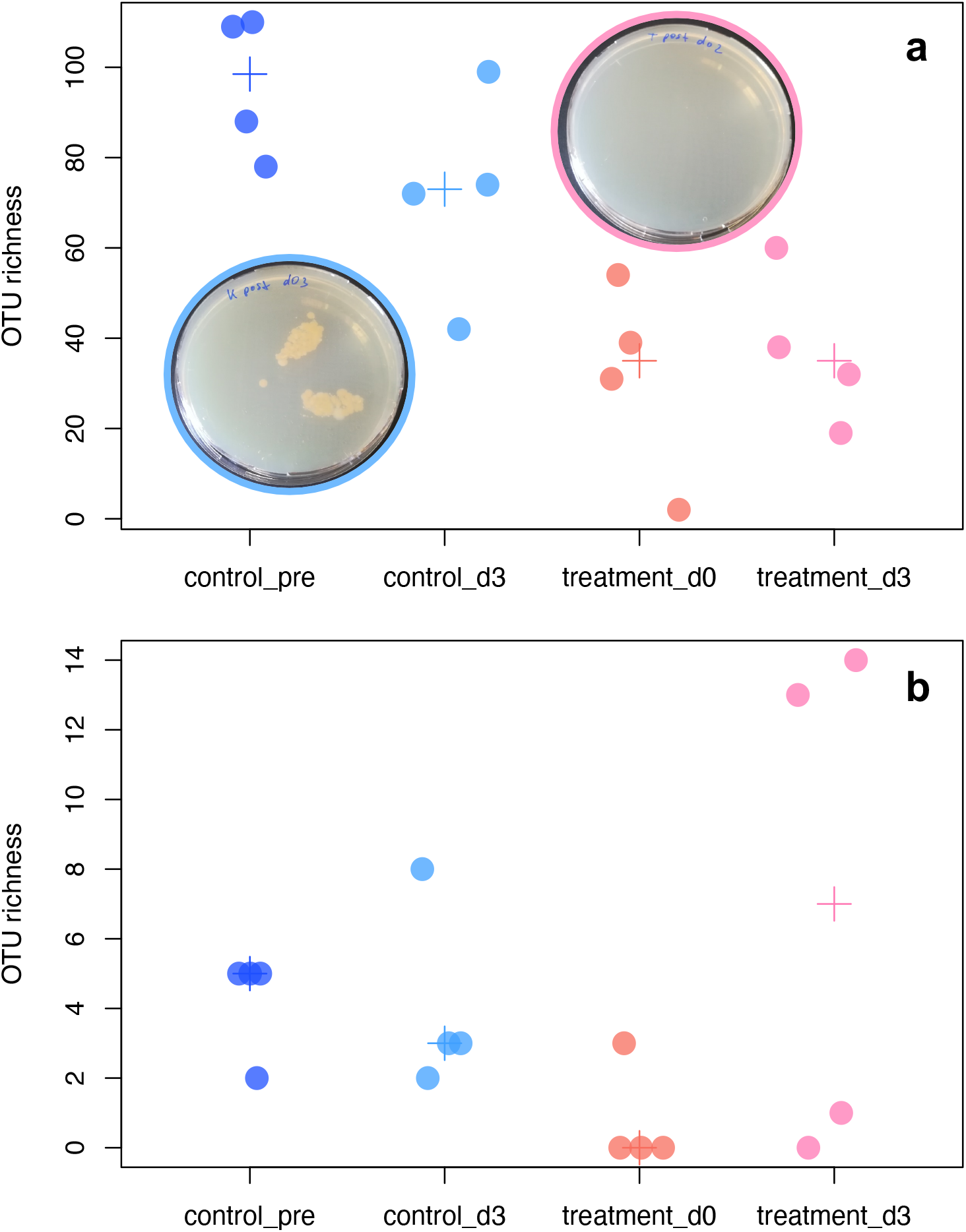
Number of OTUs (OTU richness) detected on leaf surfaces of *Arabidopsis thaliana*. Number of bacterial (**a**) and fungal (**b**) OTUs detected in the samples of control leaf surfaces prior (control_pre) and three days after the treatment (control_d3) as well as of leaf surfaces directly after PDI treatment (treatment_d0) and three days later (treatment_d3, closed circles). For each group, the median OTU number is given as a +. Leaves of control and treatment plants were placed onto LB-medium in order to observe bacterial growth on leaf surfaces. Examples for control (blue frame) and treatment (pink frame) leaves are shown in (**a**).

Next to effects on microbial richness, community composition was also strongly affected by PDI treatment. Both bacterial (Fig. 2a) and fungal (Fig. 2b) communities differed between treatments (Permutation test for Bray-Curtis distance-based redundancy analysis under reduced model: bacteria: *F*3,12 = 1.54, *p* < 0.001; fungi: *F*3,12 = 2.14, *p* < 0.01). Control plants shared a large number of bacterial OTUs whereas the treated plants shared fewer OTUs with plants in other treatment groups (see Venn diagram in Fig. 2a). Fungal OTUs were mostly unique to the plants that received different treatments, i.e. we detected little overlap of the OTUs across samples of different treatments (see Venn diagram in Fig. 2b). Additional to these community-wide changes, we analyzed the responses of individual bacterial and fungal taxa to the PDI treatment. These results suggest that individual taxa responded more strongly to the treatment than others (Supporting information S1), which, however, has to be viewed with caution due to the small sample size.

**Fig. 2.**
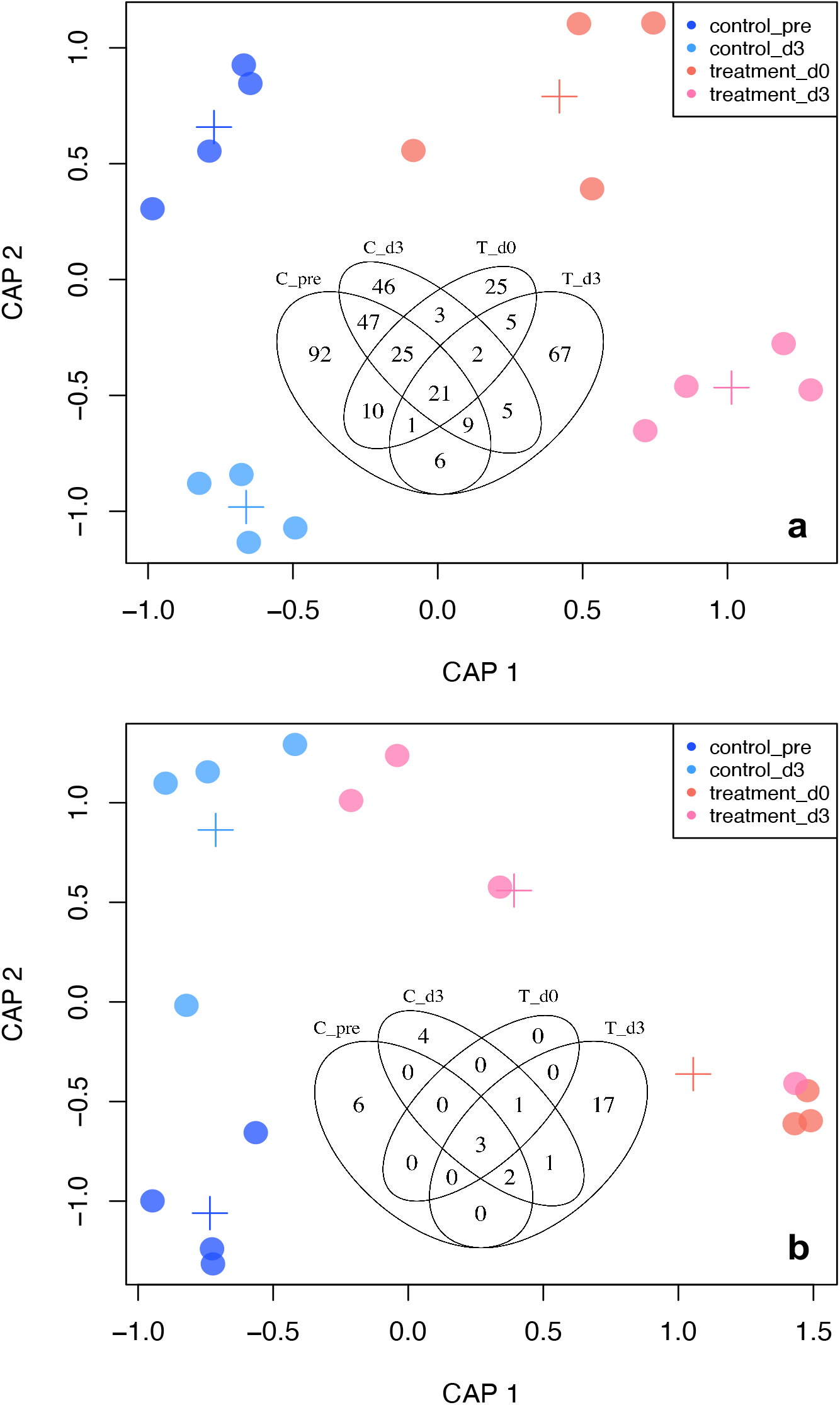
Composition of bacterial (**a**) and fungal (**b**) OUT communities associated with leaf surfaces of *Arabidopsis thaliana*. Ordination is based on Bray-Curtis distance-based redundancy analysis. The closer two closed circles the more similar is the community composition. For each treatment group (control leaf surfaces prior (control_pre) and three days after the treatment (control_d3) as well as of leaf surfaces directly after PDI treatment (treatment_d0) and three days later (treatment_d3)) the four samples are shown. Centroids of treatment groups are denoted by a +. Please note, that for the ordination visualizing the similarities in fungal communities (**b**), we jittered the coordinates of the samples to make overlapping circles visible (the circles on the bottom right completely overlapped as these are the samples from which no fungal OTUs could be detected). Venn diagrams showing the number of OTUs shared by the treatment groups are shown as insets.

Fastaq files of samples containing the sequences of the bacterial and fungal OTUs associated with *A. thaliana* are deposited at the NCBI Sequence Read Archive (SRA) under the BioProject accession PRJNA697830.

## Discussion

Our study confirmed the photoinactivation of microbes by the chlorine e6 derivative B17-0024 [9]. Additionally, we documented effects of PDI on whole bacterial and fungal communities associated with plant surfaces characterized by cultivation-independent methods. PDI strongly reduced the number of microbial OTUs, sometimes OTU-numbers even dropped to zero in fungi. On top of the effects on microbial richness, the community composition significantly changed due to the application of the photosensitizer. The bacterial communities isolated from control leaves prior to the PDI treatment or three days later shared the highest number of OTUs whereas the treated communities comprised a higher proportion of unique OTUs. Overall, beta-diversity, i.e. the difference in the inventory of OTUs between treatment groups, was higher in fungal communities indicated by few OTUs shared by two or more treatment groups. The implications of these results are thus twofold: First, PDI is a viable method to reduce the diversity of microbial communities associated with plant surfaces. Second, although it seems that some bacterial and fungal taxa are more heavily affected than others, the effect of PDI is not taxon-specific but generally affects bacteria and fungi. The latter conclusion suggests that PDI may not be suited to specifically attack pathogens but has general effects on the plant microbiome. This is a logical consequence of the mode of action of PDI: in contrast to targeted approaches, the reactive oxygen species generated by the photosensitizing agent kill all types of microorganisms, i.e. bacteria, fungi and viruses [20]. Although physico-chemical properties such as the net charge of the molecule, its size in terms of molecular weight and its lipophilicity can make uptake into or attachment to specific genera more likely, the approach is very general when compared to other antibacterial approached such as phage therapy [21]. However, due to its generalist properties PDI might initiate long lasting protective approaches for plant protection such as effective microorganisms but eradicating a pre-existing contamination with pathogens and therefore allow for controlled establishment of a protective microfauna.

Apart from implications for the application of PDI in agricultural settings, our results may stimulate microbiome research exploiting the effects of PDI. The manipulation of the plant microbiome is a challenging tasks and often involves labor-intensive preparations such as plant cultivation from surface-sterilized seeds in hermetically sealed sterile containers [4,22]. These plants can be subsequently inoculated with either individual strains or artificial communities. In such approaches the plants usually grow in highly artificial environments which may limit the relevance for natural systems. Furthermore, inoculations of field- or greenhouse-grown plants often are not persistent indicating that inoculated microbes cannot establish in their habitat, which may be inhibited by a diverse natural microbial community that occupies the available niches on plant surfaces [6]. In order to facilitate the establishment of strains or communities on plant surfaces, it may be helpful to first disrupt the community structure of resident microbial communities. For this task, environmentally friendly PDI may be a suitable way to condition plant surfaces to make them more susceptible for inoculants. In the long run, a two-step treatment of plants – plant preparation with PDI and subsequent inoculation – may increase the success of microbial application to protect plants against pathogens and to increase crop yield. To conclude, Photodynamic Inactivation of microbes on plant surfaces is a promising new tool for agricultural application and potentially also to foster academic research on the plant microbiome.

## Acknowledgement

We thank Suncor AgroScience (Mississauga Ontario, Canada) for providing the photoactive compound chlorine e6 derivative B17-0024.

