## Supplementary material for "Photodynamic Inactivation reduces the diversity and changes the composition of bacterial and fungal communities associated with leaf surfaces": S1

S1 - Bacterial OTUs detected on leaf surfaces or Arabidopis thaliana. The mean abundance of each taxon on the four treatment groups is given, as well as the p-value of a Kruskal-Wallis rank sum test testing for differences in the abundance between treatment groups

| | taxon | freq | mean_pre | mean_post_control_d3 | mean_post_treat_d0 | mean_post_treat_d3 | p-value |
|---|---|---|---|---|---|---|---|
| 1 | OTU_1957 | 4 | 6.809477019 | 0 | 0 | 0 | 0.002172664 |
| 2 | OTU_993 | 8 | 5.135204367 | 6.718807919 | 0 | 0 | 0.002747404 |
| 3 | OTU_734 | 6 | 3.391755841 | 8.03505208 | 0 | 0 | 0.010204918 |
| 4 | OTU_5264 | 3 | 2.210930722 | 0 | 0 | 0 | 0.016532305 |
| 5 | OTU_3459 | 3 | 3.924700228 | 0 | 0 | 0 | 0.016532305 |
| 6 | OTU_4947 | 3 | 3.697230967 | 0 | 0 | 0 | 0.016532305 |
| 7 | OTU_3008 | 3 | 3.795123225 | 0 | 0 | 0 | 0.016532305 |
| 8 | OTU_3762 | 3 | 4.680173508 | 0 | 0 | 0 | 0.016532305 |
| 9 | OTU_3331 | 3 | 0 | 3.973314478 | 0 | 0 | 0.016532305 |
| 10 | OTU_446 | 10 | 7.331305417 | 8.985238406 | 2.03908085 | 1.924810358 | 0.018126546 |
| 11 | OTU_819 | 7 | 2.861861228 | 6.131283078 | 1.377962662 | 0 | 0.02439948 |
| 12 | OTU_1136 | 6 | 5.605220094 | 3.43080673 | 0 | 0 | 0.031291122 |
| 13 | OTU_604 | 11 | 7.322415217 | 6.130487385 | 5.610608485 | 0 | 0.031808527 |
| 14 | OTU_1004 | 9 | 5.541021728 | 5.247387383 | 1.470483527 | 0 | 0.033165232 |
| 15 | OTU_1553 | 6 | 1.108993161 | 5.709398268 | 0 | 1.958076354 | 0.035583266 |
| 16 | OTU_2557 | 6 | 4.513091401 | 1.016098652 | 1.387830705 | 0 | 0.03750606 |
| 17 | OTU_539 | 7 | 4.085060814 | 8.20725473 | 1.802625071 | 0 | 0.037784465 |
| 18 | OTU_654 | 6 | 5.36755751 | 5.229230476 | 0 | 0 | 0.038505143 |
| 19 | OTU_3091 | 4 | 0.976050108 | 3.589870546 | 0 | 0 | 0.040038977 |
| 20 | OTU_1419 | 9 | 6.487190027 | 6.18732768 | 1.922249892 | 0 | 0.04256693 |
| 21 | OTU_949 | 7 | 3.040233706 | 6.242566738 | 1.469750225 | 0 | 0.042632768 |
| 22 | OTU_171 | 13 | 6.561037036 | 8.991394565 | 3.504821436 | 5.717836125 | 0.049595721 |
| 23 | OTU_1774 | 7 | 6.469924806 | 1.607756463 | 3.289791997 | 0 | 0.051677338 |
| 24 | OTU_3821 | 4 | 3.60263375 | 0.993016271 | 0 | 0 | 0.054463069 |
| 25 | OTU_3543 | 4 | 1.453271412 | 4.438128457 | 0 | 0 | 0.054463069 |
| 26 | OTU_1092 | 8 | 4.636883721 | 5.926549064 | 1.517144609 | 0 | 0.056755603 |
| 27 | OTU_1468 | 5 | 4.577144921 | 2.776117034 | 0 | 0 | 0.061899099 |
| 28 | OTU_2281 | 5 | 4.468994419 | 2.677815724 | 0 | 0 | 0.061899099 |
| 29 | OTU_2336 | 5 | 3.062932012 | 4.877536584 | 0 | 0 | 0.061899099 |
| 30 | OTU_1470 | 8 | 3.367958251 | 5.144305675 | 1.652270143 | 0 | 0.06385173 |
| 31 | OTU_1404 | 5 | 4.279490617 | 2.765946229 | 0 | 0 | 0.070524808 |
| 32 | OTU_1060 | 7 | 4.818214826 | 2.445428808 | 1.426768466 | 0 | 0.075690838 |
| 33 | OTU_376 | 8 | 5.122437232 | 3.873378029 | 1.559028263 | 0 | 0.082523583 |
| 34 | OTU_2032 | 5 | 3.330038786 | 0 | 2.71289774 | 0 | 0.083850507 |
| 35 | OTU_823 | 8 | 7.396959705 | 3.403376403 | 4.13040619 | 0 | 0.086988644 |
| 36 | OTU_156 | 13 | 9.942639347 | 7.601978406 | 7.314398172 | 4.13212822 | 0.087871417 |
| 37 | OTU_3804 | 4 | 3.133447673 | 0 | 1.305890741 | 0 | 0.090391026 |
| 38 | OTU_2924 | 4 | 3.99276401 | 0 | 1.803145809 | 0 | 0.090391026 |
| 39 | OTU_7726 | 2 | 1.938387447 | 0 | 0 | 0 | 0.09369079 |
| 40 | OTU_2301 | 2 | 2.7888304 | 0 | 0 | 0 | 0.09369079 |
| 41 | OTU_9567 | 2 | 2.73982926 | 0 | 0 | 0 | 0.09369079 |
| 42 | OTU_3096 | 2 | 2.443276887 | 0 | 0 | 0 | 0.09369079 |
| 43 | OTU_9756 | 2 | 2.61071996 | 0 | 0 | 0 | 0.09369079 |
| 44 | OTU_12971 | 2 | 1.624398052 | 0 | 0 | 0 | 0.09369079 |
| 45 | OTU_8595 | 2 | 2.302565682 | 0 | 0 | 0 | 0.09369079 |
| 46 | OTU_9044 | 2 | 2.250290909 | 0 | 0 | 0 | 0.09369079 |
| 47 | OTU_10945 | 2 | 1.914119687 | 0 | 0 | 0 | 0.09369079 |
| 48 | OTU_8914 | 2 | 2.270106124 | 0 | 0 | 0 | 0.09369079 |
| 49 | OTU_3140 | 2 | 2.794758842 | 0 | 0 | 0 | 0.09369079 |
| 50 | OTU_3492 | 2 | 2.536826232 | 0 | 0 | 0 | 0.09369079 |
| 51 | OTU_3789 | 2 | 2.485034508 | 0 | 0 | 0 | 0.09369079 |
| 52 | OTU_5863 | 2 | 2.859845725 | 0 | 0 | 0 | 0.09369079 |
| 53 | OTU_5791 | 2 | 1.897595406 | 0 | 0 | 0 | 0.09369079 |
| 54 | OTU_8560 | 2 | 2.934805459 | 0 | 0 | 0 | 0.09369079 |
| 55 | OTU_8485 | 2 | 2.454088672 | 0 | 0 | 0 | 0.09369079 |
| 56 | OTU_4462 | 2 | 0 | 2.868885229 | 0 | 0 | 0.09369079 |
| 57 | OTU_2712 | 2 | 0 | 2.974327087 | 0 | 0 | 0.09369079 |
| 58 | OTU_7664 | 2 | 0 | 2.432126189 | 0 | 0 | 0.09369079 |
| 59 | OTU_6913 | 2 | 0 | 0 | 0 | 3.452632682 | 0.09369079 |
| 60 | OTU_3215 | 2 | 0 | 0 | 0 | 3.795979 | 0.09369079 |
| 61 | OTU_1908 | 9 | 4.967676146 | 3.451641792 | 2.679596914 | 0 | 0.094924801 |
| 62 | OTU_1919 | 6 | 3.814011722 | 1.039542495 | 2.693428644 | 0 | 0.125479559 |
| 63 | OTU_342 | 15 | 6.250390527 | 7.830535874 | 5.83543698 | 7.27900563 | 0.155847115 |
| 64 | OTU_554 | 7 | 6.884539181 | 1.664552871 | 1.950584212 | 1.693478433 | 0.156163222 |

| | | | | | | | |
|---|---|---|---|---|---|---|---|
| 65 | OTU_1336 | 4 | 2.728228492 | 3.960832877 | 0 | 0 | 0.16994805 |
| 66 | OTU_1791 | 8 | 2.928813937 | 3.859417521 | 2.924227165 | 0 | 0.177251888 |
| 67 | OTU_5250 | 4 | 1.855319587 | 0 | 2.165792114 | 0 | 0.178418666 |
| 68 | OTU_1581 | 6 | 4.617387498 | 2.533657747 | 1.71667285 | 0 | 0.188315979 |
| 69 | OTU_2257 | 6 | 4.673166174 | 2.629395486 | 1.736548343 | 0 | 0.188315979 |
| 70 | OTU_1363 | 6 | 4.69796969 | 0 | 2.915464008 | 1.507015345 | 0.196616319 |
| 71 | OTU_3756 | 3 | 2.53684148 | 1.064825447 | 0 | 0 | 0.203162351 |
| 72 | OTU_3131 | 3 | 2.721150357 | 1.220930125 | 0 | 0 | 0.203162351 |
| 73 | OTU_2216 | 3 | 1.22199642 | 3.143039721 | 0 | 0 | 0.203162351 |
| 74 | OTU_6156 | 3 | 2.170292376 | 0.934949892 | 0 | 0 | 0.203162351 |
| 75 | OTU_3014 | 3 | 1.021187513 | 2.890048781 | 0 | 0 | 0.203162351 |
| 76 | OTU_1329 | 3 | 1.150524531 | 3.665027412 | 0 | 0 | 0.203162351 |
| 77 | OTU_1660 | 3 | 1.188027964 | 2.830530519 | 0 | 0 | 0.203162351 |
| 78 | OTU_233 | 3 | 0.841657626 | 0 | 0 | 3.845126038 | 0.203162351 |
| 79 | OTU_6186 | 3 | 1.272978081 | 2.85415464 | 0 | 0 | 0.203162351 |
| 80 | OTU_2757 | 3 | 0 | 0 | 1.766118983 | 3.918087002 | 0.203162351 |
| 81 | OTU_615 | 3 | 0 | 1.151258215 | 0 | 3.73043766 | 0.203162351 |
| 82 | OTU_809 | 8 | 4.750810858 | 3.01536892 | 0 | 3.353673628 | 0.209612165 |
| 83 | OTU_500 | 10 | 4.23430605 | 5.958008224 | 1.336468141 | 3.485556882 | 0.221277594 |
| 84 | OTU_117 | 14 | 6.63549157 | 8.767102346 | 5.349030774 | 6.78902675 | 0.222254102 |
| 85 | OTU_102 | 14 | 7.627329199 | 4.213072321 | 6.779200162 | 5.563924396 | 0.233829614 |
| 86 | OTU_106 | 15 | 7.449870884 | 4.924490375 | 8.14057723 | 5.898651658 | 0.239340118 |
| 87 | OTU_2948 | 3 | 3.170912759 | 1.550416089 | 0 | 0 | 0.242953 |
| 88 | OTU_5445 | 3 | 2.244103662 | 0 | 0 | 1.400862432 | 0.242953 |
| 89 | OTU_3590 | 3 | 3.156663664 | 1.593047922 | 0 | 0 | 0.242953 |
| 90 | OTU_5048 | 3 | 2.922448229 | 1.476014044 | 0 | 0 | 0.242953 |
| 91 | OTU_2371 | 3 | 1.22199642 | 2.477844028 | 0 | 0 | 0.242953 |
| 92 | OTU_91 | 9 | 8.895516037 | 6.152375519 | 2.640893621 | 2.391787364 | 0.249768721 |
| 93 | OTU_1410 | 6 | 2.561164238 | 3.654005931 | 1.489461386 | 0 | 0.276244692 |
| 94 | OTU_3378 | 3 | 3.241836484 | 0 | 0 | 2.077104459 | 0.278702327 |
| 95 | OTU_5746 | 3 | 2.523195835 | 1.475353629 | 0 | 0 | 0.278702327 |
| 96 | OTU_1889 | 3 | 1.529888885 | 2.924433354 | 0 | 0 | 0.278702327 |
| 97 | OTU_4905 | 3 | 0 | 2.575199728 | 1.797277083 | 0 | 0.278702327 |
| 98 | OTU_479 | 3 | 0 | 2.003377084 | 0 | 2.127578658 | 0.278702327 |
| 99 | OTU_630 | 3 | 0 | 2.642458096 | 0 | 2.211274066 | 0.278702327 |
| 100 | OTU_1259 | 11 | 5.466588399 | 3.635928156 | 4.588149935 | 1.558631024 | 0.287074383 |
| 101 | OTU_956 | 6 | 3.719920297 | 3.066111736 | 1.709884633 | 0 | 0.28803503 |
| 102 | OTU_1681 | 6 | 2.641088656 | 3.69608513 | 0 | 1.720303325 | 0.298498454 |
| 103 | OTU_4543 | 5 | 0.750667037 | 2.101934411 | 0 | 3.077723626 | 0.30558278 |
| 104 | OTU_5 | 5 | 0 | 1.188027964 | 3.640499544 | 5.286605956 | 0.30558278 |
| 105 | OTU_225 | 13 | 7.695895818 | 3.594071323 | 5.86271828 | 8.638047914 | 0.359297499 |
| 106 | OTU_1046 | 4 | 0 | 1.021187513 | 1.702196364 | 4.26734348 | 0.36623485 |
| 107 | OTU_20213 | 1 | 0.672658055 | 0 | 0 | 0 | 0.391625176 |
| 108 | OTU_16241 | 1 | 0.916443363 | 0 | 0 | 0 | 0.391625176 |
| 109 | OTU_10852 | 1 | 1.468682761 | 0 | 0 | 0 | 0.391625176 |
| 110 | OTU_15771 | 1 | 0.958227435 | 0 | 0 | 0 | 0.391625176 |
| 111 | OTU_1718 | 1 | 1.542117957 | 0 | 0 | 0 | 0.391625176 |
| 112 | OTU_12254 | 1 | 1.163295394 | 0 | 0 | 0 | 0.391625176 |
| 113 | OTU_18814 | 1 | 0.784139409 | 0 | 0 | 0 | 0.391625176 |
| 114 | OTU_20686 | 1 | 0.750667037 | 0 | 0 | 0 | 0.391625176 |
| 115 | OTU_17804 | 1 | 1.205425168 | 0 | 0 | 0 | 0.391625176 |
| 116 | OTU_11120 | 1 | 1.102602304 | 0 | 0 | 0 | 0.391625176 |
| 117 | OTU_4243 | 1 | 1.658093691 | 0 | 0 | 0 | 0.391625176 |
| 118 | OTU_10080 | 1 | 1.50075033 | 0 | 0 | 0 | 0.391625176 |
| 119 | OTU_10399 | 1 | 1.184975071 | 0 | 0 | 0 | 0.391625176 |
| 120 | OTU_8528 | 1 | 1.243144513 | 0 | 0 | 0 | 0.391625176 |
| 121 | OTU_7339 | 1 | 1.184975071 | 0 | 0 | 0 | 0.391625176 |
| 122 | OTU_3641 | 1 | 1.458971372 | 0 | 0 | 0 | 0.391625176 |
| 123 | OTU_19640 | 1 | 1.11558552 | 0 | 0 | 0 | 0.391625176 |
| 124 | OTU_10258 | 1 | 1.163295394 | 0 | 0 | 0 | 0.391625176 |
| 125 | OTU_17806 | 1 | 0.937940147 | 0 | 0 | 0 | 0.391625176 |
| 126 | OTU_12037 | 1 | 1.075143646 | 0 | 0 | 0 | 0.391625176 |
| 127 | OTU_4411 | 1 | 1.473442127 | 0 | 0 | 0 | 0.391625176 |
| 128 | OTU_19553 | 1 | 0.784139409 | 0 | 0 | 0 | 0.391625176 |
| 129 | OTU_20376 | 1 | 1.075143646 | 0 | 0 | 0 | 0.391625176 |
| 130 | OTU_14431 | 1 | 1.11558552 | 0 | 0 | 0 | 0.391625176 |
| 131 | OTU_13183 | 1 | 1.151946449 | 0 | 0 | 0 | 0.391625176 |
| 132 | OTU_20544 | 1 | 1.060588398 | 0 | 0 | 0 | 0.391625176 |
| 133 | OTU_11624 | 1 | 1.19534504 | 0 | 0 | 0 | 0.391625176 |
| 134 | OTU_10806 | 1 | 1.160272273 | 0 | 0 | 0 | 0.391625176 |
| 135 | OTU_15661 | 1 | 0.885421326 | 0 | 0 | 0 | 0.391625176 |
| 136 | OTU_11194 | 1 | 1.04841802 | 0 | 0 | 0 | 0.391625176 |

| | | | | | | | |
|---|---|---|---|---|---|---|---|
| 137 | OTU_20303 | 1 | 1.021187513 | 0 | 0 | 0 | 0.391625176 |
| 138 | OTU_14714 | 1 | 0.924444786 | 0 | 0 | 0 | 0.391625176 |
| 139 | OTU_10633 | 1 | 1.085758556 | 0 | 0 | 0 | 0.391625176 |
| 140 | OTU_11523 | 1 | 1.188027964 | 0 | 0 | 0 | 0.391625176 |
| 141 | OTU_12643 | 1 | 0.991732001 | 0 | 0 | 0 | 0.391625176 |
| 142 | OTU_13711 | 1 | 0.959655259 | 0 | 0 | 0 | 0.391625176 |
| 143 | OTU_16550 | 1 | 0.864202849 | 0 | 0 | 0 | 0.391625176 |
| 144 | OTU_19498 | 1 | 0.734023313 | 0 | 0 | 0 | 0.391625176 |
| 145 | OTU_18792 | 1 | 0.764089346 | 0 | 0 | 0 | 0.391625176 |
| 146 | OTU_5336 | 1 | 1.260399761 | 0 | 0 | 0 | 0.391625176 |
| 147 | OTU_14124 | 1 | 0.864202849 | 0 | 0 | 0 | 0.391625176 |
| 148 | OTU_8168 | 1 | 1.42208381 | 0 | 0 | 0 | 0.391625176 |
| 149 | OTU_7661 | 1 | 1.755579284 | 0 | 0 | 0 | 0.391625176 |
| 150 | OTU_16404 | 1 | 0.96940245 | 0 | 0 | 0 | 0.391625176 |
| 151 | OTU_17668 | 1 | 1.234810939 | 0 | 0 | 0 | 0.391625176 |
| 152 | OTU_12544 | 1 | 1.435443697 | 0 | 0 | 0 | 0.391625176 |
| 153 | OTU_19854 | 1 | 1.107369599 | 0 | 0 | 0 | 0.391625176 |
| 154 | OTU_9678 | 1 | 1.296348981 | 0 | 0 | 0 | 0.391625176 |
| 155 | OTU_14379 | 1 | 1.19212162 | 0 | 0 | 0 | 0.391625176 |
| 156 | OTU_20305 | 1 | 0.792115084 | 0 | 0 | 0 | 0.391625176 |
| 157 | OTU_13411 | 1 | 1.24798722 | 0 | 0 | 0 | 0.391625176 |
| 158 | OTU_14237 | 1 | 1.234810939 | 0 | 0 | 0 | 0.391625176 |
| 159 | OTU_21557 | 1 | 0.655808129 | 0 | 0 | 0 | 0.391625176 |
| 160 | OTU_19578 | 1 | 0.724388183 | 0 | 0 | 0 | 0.391625176 |
| 161 | OTU_21739 | 1 | 0.571052107 | 0 | 0 | 0 | 0.391625176 |
| 162 | OTU_17231 | 1 | 1.12987736 | 0 | 0 | 0 | 0.391625176 |
| 163 | OTU_17256 | 1 | 1.401772909 | 0 | 0 | 0 | 0.391625176 |
| 164 | OTU_1991 | 1 | 2.28296424 | 0 | 0 | 0 | 0.391625176 |
| 165 | OTU_19560 | 1 | 1.063204502 | 0 | 0 | 0 | 0.391625176 |
| 166 | OTU_17686 | 1 | 1.03792805 | 0 | 0 | 0 | 0.391625176 |
| 167 | OTU_16514 | 1 | 0.875282727 | 0 | 0 | 0 | 0.391625176 |
| 168 | OTU_11639 | 1 | 1.234776305 | 0 | 0 | 0 | 0.391625176 |
| 169 | OTU_18529 | 1 | 0.914207122 | 0 | 0 | 0 | 0.391625176 |
| 170 | OTU_11020 | 1 | 1.590662718 | 0 | 0 | 0 | 0.391625176 |
| 171 | OTU_16652 | 1 | 1.086824971 | 0 | 0 | 0 | 0.391625176 |
| 172 | OTU_20919 | 1 | 0.949336937 | 0 | 0 | 0 | 0.391625176 |
| 173 | OTU_16974 | 1 | 1.361403912 | 0 | 0 | 0 | 0.391625176 |
| 174 | OTU_4930 | 1 | 1.522378638 | 0 | 0 | 0 | 0.391625176 |
| 175 | OTU_17301 | 1 | 1.12987736 | 0 | 0 | 0 | 0.391625176 |
| 176 | OTU_4 | 1 | 0 | 0 | 1.254491232 | 0 | 0.391625176 |
| 177 | OTU_311 | 1 | 0 | 0 | 2.761117504 | 0 | 0.391625176 |
| 178 | OTU_21322 | 1 | 0 | 0 | 1.340955469 | 0 | 0.391625176 |
| 179 | OTU_21682 | 1 | 0 | 0 | 1.299567407 | 0 | 0.391625176 |
| 180 | OTU_20014 | 1 | 0 | 0 | 1.470879022 | 0 | 0.391625176 |
| 181 | OTU_14934 | 1 | 0 | 0 | 1.929969426 | 0 | 0.391625176 |
| 182 | OTU_19074 | 1 | 0 | 0 | 1.521676231 | 0 | 0.391625176 |
| 183 | OTU_21440 | 1 | 0 | 0 | 1.340955469 | 0 | 0.391625176 |
| 184 | OTU_14255 | 1 | 0 | 0 | 1.497171172 | 0 | 0.391625176 |
| 185 | OTU_20917 | 1 | 0 | 0 | 1.411736093 | 0 | 0.391625176 |
| 186 | OTU_16269 | 1 | 0 | 0 | 1.880960489 | 0 | 0.391625176 |
| 187 | OTU_21315 | 1 | 0 | 0 | 1.340955469 | 0 | 0.391625176 |
| 188 | OTU_21397 | 1 | 0 | 0 | 1.340955469 | 0 | 0.391625176 |
| 189 | OTU_20330 | 1 | 0 | 0 | 1.442518476 | 0 | 0.391625176 |
| 190 | OTU_17877 | 1 | 0 | 0 | 1.544621754 | 0 | 0.391625176 |
| 191 | OTU_21629 | 1 | 0 | 0 | 1.299567407 | 0 | 0.391625176 |
| 192 | OTU_3336 | 1 | 0 | 0 | 0.957542036 | 0 | 0.391625176 |
| 193 | OTU_17167 | 1 | 0 | 0 | 1.095229291 | 0 | 0.391625176 |
| 194 | OTU_13258 | 1 | 0 | 0 | 1.377962662 | 0 | 0.391625176 |
| 195 | OTU_11475 | 1 | 0 | 0 | 1.350836652 | 0 | 0.391625176 |
| 196 | OTU_16243 | 1 | 0 | 0 | 1.272478878 | 0 | 0.391625176 |
| 197 | OTU_9704 | 1 | 0 | 0 | 1.336468141 | 0 | 0.391625176 |
| 198 | OTU_20721 | 1 | 0 | 0 | 0.802926831 | 0 | 0.391625176 |
| 199 | OTU_15750 | 1 | 0 | 0 | 1.28957157 | 0 | 0.391625176 |
| 200 | OTU_6000 | 1 | 0 | 0 | 1.764447595 | 0 | 0.391625176 |
| 201 | OTU_20077 | 1 | 0 | 1.138215837 | 0 | 0 | 0.391625176 |
| 202 | OTU_382 | 1 | 0 | 1.65294829 | 0 | 0 | 0.391625176 |
| 203 | OTU_15175 | 1 | 0 | 0.903263702 | 0 | 0 | 0.391625176 |
| 204 | OTU_14756 | 1 | 0 | 0.927880865 | 0 | 0 | 0.391625176 |
| 205 | OTU_14377 | 1 | 0 | 1.095995133 | 0 | 0 | 0.391625176 |
| 206 | OTU_20947 | 1 | 0 | 0.783503394 | 0 | 0 | 0.391625176 |
| 207 | OTU_16056 | 1 | 0 | 1.012352721 | 0 | 0 | 0.391625176 |
| 208 | OTU_9126 | 1 | 0 | 1.048168576 | 0 | 0 | 0.391625176 |

| | | | | | | | |
|---|---|---|---|---|---|---|---|
| 209 | OTU_2728 | 1 | 0 | 1.537480683 | 0 | 0 | 0.391625176 |
| 210 | OTU_10804 | 1 | 0 | 1.251293314 | 0 | 0 | 0.391625176 |
| 211 | OTU_9291 | 1 | 0 | 1.313545348 | 0 | 0 | 0.391625176 |
| 212 | OTU_4837 | 1 | 0 | 1.840164593 | 0 | 0 | 0.391625176 |
| 213 | OTU_8867 | 1 | 0 | 1.344821188 | 0 | 0 | 0.391625176 |
| 214 | OTU_3744 | 1 | 0 | 1.639967994 | 0 | 0 | 0.391625176 |
| 215 | OTU_17512 | 1 | 0 | 1.012352721 | 0 | 0 | 0.391625176 |
| 216 | OTU_364 | 1 | 0 | 1.732692 | 0 | 0 | 0.391625176 |
| 217 | OTU_1572 | 1 | 0 | 1.972117409 | 0 | 0 | 0.391625176 |
| 218 | OTU_3853 | 1 | 0 | 1.677586272 | 0 | 0 | 0.391625176 |
| 219 | OTU_3 | 1 | 0 | 1.425067963 | 0 | 0 | 0.391625176 |
| 220 | OTU_14625 | 1 | 0 | 1.039542495 | 0 | 0 | 0.391625176 |
| 221 | OTU_17607 | 1 | 0 | 1.08230106 | 0 | 0 | 0.391625176 |
| 222 | OTU_11011 | 1 | 0 | 1.333408194 | 0 | 0 | 0.391625176 |
| 223 | OTU_4420 | 1 | 0 | 1.101918389 | 0 | 0 | 0.391625176 |
| 224 | OTU_8942 | 1 | 0 | 1.333408194 | 0 | 0 | 0.391625176 |
| 225 | OTU_9409 | 1 | 0 | 1.312727623 | 0 | 0 | 0.391625176 |
| 226 | OTU_10664 | 1 | 0 | 1.352966998 | 0 | 0 | 0.391625176 |
| 227 | OTU_5548 | 1 | 0 | 0.903263702 | 0 | 0 | 0.391625176 |
| 228 | OTU_10810 | 1 | 0 | 1.061555045 | 0 | 0 | 0.391625176 |
| 229 | OTU_6456 | 1 | 0 | 1.429832783 | 0 | 0 | 0.391625176 |
| 230 | OTU_20797 | 1 | 0 | 0.830076544 | 0 | 0 | 0.391625176 |
| 231 | OTU_35 | 1 | 0 | 0.865952847 | 0 | 0 | 0.391625176 |
| 232 | OTU_5311 | 1 | 0 | 1.473876723 | 0 | 0 | 0.391625176 |
| 233 | OTU_3111 | 1 | 0 | 1.17626794 | 0 | 0 | 0.391625176 |
| 234 | OTU_8087 | 1 | 0 | 1.869454793 | 0 | 0 | 0.391625176 |
| 235 | OTU_7179 | 1 | 0 | 1.424403174 | 0 | 0 | 0.391625176 |
| 236 | OTU_11733 | 1 | 0 | 1.320648423 | 0 | 0 | 0.391625176 |
| 237 | OTU_9295 | 1 | 0 | 1.387941933 | 0 | 0 | 0.391625176 |
| 238 | OTU_15291 | 1 | 0 | 1.130201223 | 0 | 0 | 0.391625176 |
| 239 | OTU_7459 | 1 | 0 | 1.271166299 | 0 | 0 | 0.391625176 |
| 240 | OTU_375 | 1 | 0 | 1.160272273 | 0 | 0 | 0.391625176 |
| 241 | OTU_3443 | 1 | 0 | 1.63793426 | 0 | 0 | 0.391625176 |
| 242 | OTU_15581 | 1 | 0 | 1.130201223 | 0 | 0 | 0.391625176 |
| 243 | OTU_1194 | 1 | 0 | 0 | 0 | 1.705521704 | 0.391625176 |
| 244 | OTU_12095 | 1 | 0 | 0 | 0 | 1.495742668 | 0.391625176 |
| 245 | OTU_2030 | 1 | 0 | 0 | 0 | 1.829657253 | 0.391625176 |
| 246 | OTU_3090 | 1 | 0 | 0 | 0 | 1.7486054 | 0.391625176 |
| 247 | OTU_975 | 1 | 0 | 0 | 0 | 1.538862048 | 0.391625176 |
| 248 | OTU_3074 | 1 | 0 | 0 | 0 | 1.910594493 | 0.391625176 |
| 249 | OTU_7007 | 1 | 0 | 0 | 0 | 1.731149474 | 0.391625176 |
| 250 | OTU_5808 | 1 | 0 | 0 | 0 | 1.419523634 | 0.391625176 |
| 251 | OTU_18819 | 1 | 0 | 0 | 0 | 1.239948124 | 0.391625176 |
| 252 | OTU_2517 | 1 | 0 | 0 | 0 | 1.742880118 | 0.391625176 |
| 253 | OTU_6917 | 1 | 0 | 0 | 0 | 1.357978224 | 0.391625176 |
| 254 | OTU_14075 | 1 | 0 | 0 | 0 | 1.419523634 | 0.391625176 |
| 255 | OTU_16939 | 1 | 0 | 0 | 0 | 1.28877574 | 0.391625176 |
| 256 | OTU_18188 | 1 | 0 | 0 | 0 | 1.242057605 | 0.391625176 |
| 257 | OTU_10508 | 1 | 0 | 0 | 0 | 1.613132743 | 0.391625176 |
| 258 | OTU_14729 | 1 | 0 | 0 | 0 | 1.367227626 | 0.391625176 |
| 259 | OTU_21036 | 1 | 0 | 0 | 0 | 1.048758482 | 0.391625176 |
| 260 | OTU_2122 | 1 | 0 | 0 | 0 | 1.861067734 | 0.391625176 |
| 261 | OTU_19644 | 1 | 0 | 0 | 0 | 1.18837474 | 0.391625176 |
| 262 | OTU_19458 | 1 | 0 | 0 | 0 | 1.18837474 | 0.391625176 |
| 263 | OTU_17650 | 1 | 0 | 0 | 0 | 1.242057605 | 0.391625176 |
| 264 | OTU_20217 | 1 | 0 | 0 | 0 | 1.125280503 | 0.391625176 |
| 265 | OTU_8546 | 1 | 0 | 0 | 0 | 1.833116926 | 0.391625176 |
| 266 | OTU_20288 | 1 | 0 | 0 | 0 | 1.125280503 | 0.391625176 |
| 267 | OTU_8944 | 1 | 0 | 0 | 0 | 1.575589259 | 0.391625176 |
| 268 | OTU_18341 | 1 | 0 | 0 | 0 | 1.594849281 | 0.391625176 |
| 269 | OTU_19502 | 1 | 0 | 0 | 0 | 1.18837474 | 0.391625176 |
| 270 | OTU_21047 | 1 | 0 | 0 | 0 | 1.048758482 | 0.391625176 |
| 271 | OTU_18837 | 1 | 0 | 0 | 0 | 1.18837474 | 0.391625176 |
| 272 | OTU_20935 | 1 | 0 | 0 | 0 | 1.048758482 | 0.391625176 |
| 273 | OTU_21174 | 1 | 0 | 0 | 0 | 1.048758482 | 0.391625176 |
| 274 | OTU_21044 | 1 | 0 | 0 | 0 | 1.048758482 | 0.391625176 |
| 275 | OTU_9029 | 1 | 0 | 0 | 0 | 1.802816694 | 0.391625176 |
| 276 | OTU_21654 | 1 | 0 | 0 | 0 | 1.048758482 | 0.391625176 |
| 277 | OTU_18020 | 1 | 0 | 0 | 0 | 1.533678806 | 0.391625176 |
| 278 | OTU_20931 | 1 | 0 | 0 | 0 | 1.048758482 | 0.391625176 |
| 279 | OTU_4231 | 1 | 0 | 0 | 0 | 1.968770278 | 0.391625176 |
| 280 | OTU_21354 | 1 | 0 | 0 | 0 | 1.048758482 | 0.391625176 |

| | | | | | | | |
|---|---|---|---|---|---|---|---|
| 281 | OTU_18320 | 1 | 0 | 0 | 0 | 1.555242442 | 0.391625176 |
| 282 | OTU_20196 | 1 | 0 | 0 | 0 | 1.125280503 | 0.391625176 |
| 283 | OTU_12691 | 1 | 0 | 0 | 0 | 1.663003649 | 0.391625176 |
| 284 | OTU_19689 | 1 | 0 | 0 | 0 | 1.18837474 | 0.391625176 |
| 285 | OTU_15791 | 1 | 0 | 0 | 0 | 1.533678806 | 0.391625176 |
| 286 | OTU_16646 | 1 | 0 | 0 | 0 | 1.594849281 | 0.391625176 |
| 287 | OTU_18501 | 1 | 0 | 0 | 0 | 1.242057605 | 0.391625176 |
| 288 | OTU_9353 | 1 | 0 | 0 | 0 | 1.887007387 | 0.391625176 |
| 289 | OTU_17685 | 1 | 0 | 0 | 0 | 1.242057605 | 0.391625176 |
| 290 | OTU_21618 | 1 | 0 | 0 | 0 | 1.048758482 | 0.391625176 |
| 291 | OTU_13522 | 1 | 0 | 0 | 0 | 1.869923198 | 0.391625176 |
| 292 | OTU_14159 | 1 | 0 | 0 | 0 | 1.431626425 | 0.391625176 |
| 293 | OTU_11337 | 1 | 0 | 0 | 0 | 1.486250095 | 0.391625176 |
| 294 | OTU_1400 | 1 | 0 | 0 | 0 | 2.015705069 | 0.391625176 |
| 295 | OTU_17336 | 1 | 0 | 0 | 0 | 1.556099279 | 0.391625176 |
| 296 | OTU_21296 | 1 | 0 | 0 | 0 | 1.106428658 | 0.391625176 |
| 297 | OTU_4681 | 1 | 0 | 0 | 0 | 1.575345937 | 0.391625176 |
| 298 | OTU_96 | 1 | 0 | 0 | 0 | 1.761501442 | 0.391625176 |
| 299 | OTU_41 | 1 | 0 | 0 | 0 | 1.466832346 | 0.391625176 |
| 300 | OTU_603 | 1 | 0 | 0 | 0 | 1.575345937 | 0.391625176 |
| 301 | OTU_17095 | 1 | 0 | 0 | 0 | 1.310891052 | 0.391625176 |
| 302 | OTU_5271 | 1 | 0 | 0 | 0 | 1.310891052 | 0.391625176 |
| 303 | OTU_2662 | 1 | 0 | 0 | 0 | 1.726278925 | 0.391625176 |
| 304 | OTU_974 | 1 | 0 | 0 | 0 | 2.488841017 | 0.391625176 |
| 305 | OTU_2759 | 1 | 0 | 0 | 0 | 2.24801983 | 0.391625176 |
| 306 | OTU_2705 | 1 | 0 | 0 | 0 | 1.726278925 | 0.391625176 |
| 307 | OTU_27 | 1 | 0 | 0 | 0 | 1.34793855 | 0.391625176 |
| 308 | OTU_482 | 8 | 2.736271795 | 5.182112453 | 1.254535672 | 4.78410472 | 0.401230132 |
| 309 | OTU_97 | 15 | 7.790994793 | 8.628243561 | 7.356109759 | 9.439689311 | 0.416050161 |
| 310 | OTU_4339 | 4 | 2.386829923 | 0.950924644 | 0 | 1.215891085 | 0.422227176 |
| 311 | OTU_938 | 9 | 3.676075243 | 4.476761353 | 1.912762923 | 1.303769936 | 0.429390046 |
| 312 | OTU_2276 | 5 | 2.536918908 | 2.340414401 | 1.746089902 | 0 | 0.461697243 |
| 313 | OTU_1416 | 4 | 1.140506006 | 2.635961333 | 0 | 2.287499998 | 0.471290945 |
| 314 | OTU_5066 | 4 | 1.087737317 | 2.348686391 | 1.254491232 | 0 | 0.471290945 |
| 315 | OTU_2967 | 4 | 1.160272273 | 2.669055678 | 1.521676231 | 0 | 0.485117062 |
| 316 | OTU_1685 | 6 | 2.911095178 | 0.738156685 | 1.442518476 | 2.012556396 | 0.504019229 |
| 317 | OTU_2208 | 4 | 1.205425168 | 2.474783691 | 0 | 1.759790325 | 0.53222338 |
| 318 | OTU_2792 | 4 | 1.650558075 | 0.791840764 | 0 | 1.948245098 | 0.53222338 |
| 319 | OTU_2310 | 2 | 1.376370516 | 1.11062478 | 0 | 0 | 0.541863834 |
| 320 | OTU_2886 | 2 | 1.163295394 | 1.519478581 | 0 | 0 | 0.541863834 |
| 321 | OTU_9086 | 2 | 0.937940147 | 0 | 0 | 1.190114027 | 0.541863834 |
| 322 | OTU_9073 | 2 | 1.163295394 | 1.371519494 | 0 | 0 | 0.541863834 |
| 323 | OTU_6544 | 2 | 0.995669494 | 1.064825447 | 0 | 0 | 0.541863834 |
| 324 | OTU_7853 | 2 | 1.151946449 | 1.080746899 | 0 | 0 | 0.541863834 |
| 325 | OTU_525 | 2 | 1.013027112 | 1.048168576 | 0 | 0 | 0.541863834 |
| 326 | OTU_1132 | 2 | 1.443895674 | 1.18777389 | 0 | 0 | 0.541863834 |
| 327 | OTU_6537 | 2 | 1.013027112 | 1.329522172 | 0 | 0 | 0.541863834 |
| 328 | OTU_7888 | 2 | 1.163295394 | 1.138215837 | 0 | 0 | 0.541863834 |
| 329 | OTU_4172 | 2 | 0.958227435 | 1.51483407 | 0 | 0 | 0.541863834 |
| 330 | OTU_1412 | 2 | 0.893583673 | 1.138215837 | 0 | 0 | 0.541863834 |
| 331 | OTU_4605 | 2 | 1.045420971 | 1.270202367 | 0 | 0 | 0.541863834 |
| 332 | OTU_6824 | 2 | 1.42208381 | 1.48001711 | 0 | 0 | 0.541863834 |
| 333 | OTU_1918 | 2 | 1.108663992 | 0 | 0 | 1.783206568 | 0.541863834 |
| 334 | OTU_7214 | 2 | 1.22199642 | 1.586151912 | 0 | 0 | 0.541863834 |
| 335 | OTU_6510 | 2 | 1.130201223 | 0 | 1.566851447 | 0 | 0.541863834 |
| 336 | OTU_8790 | 2 | 1.00676037 | 1.099226355 | 0 | 0 | 0.541863834 |
| 337 | OTU_2927 | 2 | 1.366875627 | 1.535720372 | 0 | 0 | 0.541863834 |
| 338 | OTU_5616 | 2 | 1.253039267 | 1.255155264 | 0 | 0 | 0.541863834 |
| 339 | OTU_5466 | 2 | 1.411589332 | 1.163845379 | 0 | 0 | 0.541863834 |
| 340 | OTU_7675 | 2 | 1.107369599 | 0 | 1.442518476 | 0 | 0.541863834 |
| 341 | OTU_6531 | 2 | 1.19212162 | 0 | 0 | 1.239948124 | 0.541863834 |
| 342 | OTU_9543 | 2 | 1.160568572 | 0 | 1.226548368 | 0 | 0.541863834 |
| 343 | OTU_8343 | 2 | 1.318297214 | 1.030705042 | 0 | 0 | 0.541863834 |
| 344 | OTU_3869 | 2 | 1.010745793 | 1.481975532 | 0 | 0 | 0.541863834 |
| 345 | OTU_8602 | 2 | 0.949336937 | 0 | 1.340955469 | 0 | 0.541863834 |
| 346 | OTU_6541 | 2 | 1.392096305 | 1.176008037 | 0 | 0 | 0.541863834 |
| 347 | OTU_8181 | 2 | 1.168334394 | 0 | 1.305890741 | 0 | 0.541863834 |
| 348 | OTU_10702 | 2 | 1.234776305 | 0 | 1.122764501 | 0 | 0.541863834 |
| 349 | OTU_98 | 2 | 0 | 0 | 1.470483527 | 1.850698233 | 0.541863834 |
| 350 | OTU_7131 | 2 | 0 | 1.201143865 | 1.484962222 | 0 | 0.541863834 |
| 351 | OTU_1980 | 2 | 0 | 0 | 1.163187433 | 1.582456942 | 0.541863834 |
| 352 | OTU_5497 | 2 | 0 | 0 | 1.986413212 | 1.658650697 | 0.541863834 |

| | | | | | | | |
|---|---|---|---|---|---|---|---|
| 353 | OTU_7800 | 2 | 0 | 0 | 1.780753206 | 1.627597821 | 0.541863834 |
| 354 | OTU_7580 | 2 | 0 | 1.251293314 | 1.254535672 | 0 | 0.541863834 |
| 355 | OTU_8364 | 2 | 0 | 0.87684245 | 0 | 1.558631024 | 0.541863834 |
| 356 | OTU_4390 | 2 | 0 | 1.08230106 | 0 | 2.099221369 | 0.541863834 |
| 357 | OTU_109 | 14 | 6.067141562 | 7.202559267 | 6.768290831 | 6.505455994 | 0.599342556 |
| 358 | OTU_968 | 10 | 5.38916073 | 4.781450584 | 3.214247703 | 1.808483301 | 0.752167778 |
| 359 | OTU_1080 | 3 | 0.893583673 | 1.064825447 | 1.194634693 | 0 | 0.756193809 |
| 360 | OTU_983 | 3 | 1.704330497 | 1.778238545 | 2.076746088 | 0 | 0.756193809 |
| 361 | OTU_2152 | 3 | 1.515478882 | 1.493591657 | 0 | 1.76525535 | 0.756193809 |
| 362 | OTU_1166 | 9 | 4.022885042 | 4.051298626 | 3.261294604 | 1.705521704 | 0.839554132 |
| 363 | OTU_507 | 11 | 4.003196808 | 3.792857809 | 2.838367036 | 3.92031812 | 0.92385536 |
| 364 | OTU_2802 | 6 | 2.206386238 | 2.127664838 | 1.617322155 | 1.472081897 | 0.970836858 |
